# A computational model for lipid-anchored polysaccharide export by the outer-membrane protein GfcD

**DOI:** 10.1101/2023.12.04.565983

**Authors:** Cecilia Fruet, Mikel Martinez-Goikoetxea, Felipe Merino, Andrei N. Lupas

**Affiliations:** Department of Protein Evolution, Max Planck Institute for Biology Tübingen, D-72076 Tübingen, Germany; Institute of Bioengineering, School of Life Sciences, École Polytechnique Fédérale de Lausanne (EPFL), CH-1015 Lausanne, Switzerland; SIB Swiss Institute of Bioinformatics, CH-1015 Lausanne, Switzerland; Cube Biotech GmbH, D-40789 Monheim, Germany

## Abstract

Many bacteria are protected by different types of polysaccharide capsules, structures formed of long repetitive glycan chains that are sometimes free and sometimes anchored to the outer membrane via lipid tails. One type, called group 4 capsule, results from expression of the *gfcABCDE-etp-etk* operon in *Escherichia coli*. Of the proteins encoded in this operon, GfcE is thought to provide the export pore for free polysaccharide chains, but none of the proteins has been implicated in the export of chains carrying a lipid anchor. For this function, GfcD has been a focus of attention as the only outer-membrane β-barrel encoded in the operon. AlphaFold predicts two β-barrel domains in GfcD, a canonical N-terminal one of 12 strands and an unusual C-terminal one of 13 strands, which features a large lateral aperture between strands β1 and β13. This immediately suggests a lateral exit gate for hydrophobic molecules into the membrane, analogous to the one proposed for the lipopolysaccharide export pore LptD. Here, we report an unsteered molecular dynamics study of GfcD embedded in the bacterial outer membrane, with the common polysaccharide anchor, lipid A, inserted in the pore of the C-terminal barrel. Our results show that the lateral aperture does not collapse during simulations, that membrane lipids nevertheless do not penetrate the barrel, but that the lipid chains of the lipid A molecule readily exit into the membrane.

**Statement of Significance:** Despite the essential role polysaccharide capsules play in the resilience of bacteria to hostile environments, many aspects of their biogenesis are still poorly understood. One aspect concerns the export of capsular polysaccharides carrying a lipid anchor, for which even the proteins mediating the process are unknown. Here we propose that one of the largest families of β-barrel proteins in the bacterial outer membrane is a key agent of this process and show by biophysical simulation that it allows the exit of lipid anchors into the membrane through a lateral opening. More generally, our model illuminates the lateral exit mechanisms proposed for the export of hydrophobic macromolecules into the bacterial outer membrane.

**Graphical abstract:** 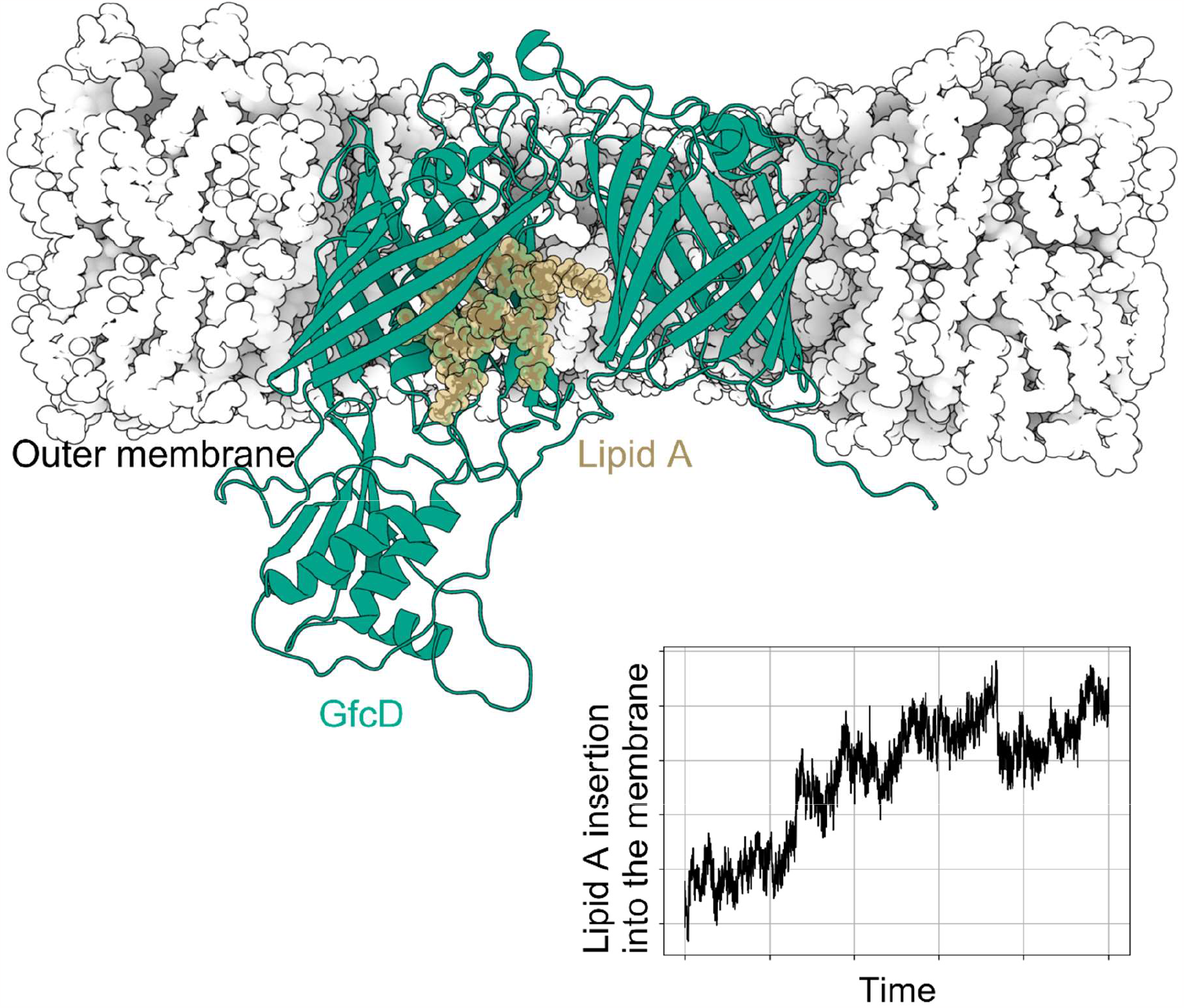

## Introduction

The bacterial capsule is a cellular structure that sheathes many bacteria, both Gram-positive and Gram-negative (1). It protects cells against changing and often hostile environments, for example against host immunity during infection, and assists in biofilm formation and cell adhesion (2). The capsule is composed of capsular polysaccharides (CPSs), long repetitive chains of several hundred identical oligosaccharide units, differing in their composition and linkage between different capsules. Although their attachment to the bacterial cell surface has not been completely elucidated, it is understood that some CPSs feature a lipid tail that anchors them to the outer leaflet of the outer membrane (3–5).

In *Escherichia coli*, there is considerable structural diversity in the CPS oligosaccharide repeat units, but there are only four known biosynthetic pathways for capsule assembly, named groups 1 to 4. The mechanism of capsule biogenesis is only well understood for group 1 (4,5), but some aspects of group 4 can be inferred from the homology of the operons governing the biosynthesis of groups 1 and 4 (4,5). Group 1 capsule production is governed by the *wzabc* operon (3–5), where Wza is an octameric translocation channel through the outer membrane, Wzc the corresponding octameric channel through the inner membrane, and Wzb a phosphatase regulating the activity of Wzc. Group 4 capsule production is driven by the *gfcABCDE-etp-etk* operon, of which the last three genes are homologs of *wzabc* (3). The first four genes of the operon, necessary for group 4 capsule expression (6), are homologous to yet another operon of *E. coli, yjbEFGH*, which is involved in the secretion of an as yet uncharacterized exopolysaccharide (3,7–10). Additionally, the homologs of *gfcBCD* in the Gram-negative bacterium *Vibrio anguillarium* are also involved in exopolysaccharide biosynthesis and transport (3,11).

Two of the four proteins encoded in this part of the operon, GfcB and GfcC, are of known structure but unknown activity, while the remaining two, GfcA and GfcD, are entirely uncharacterized at this point. During a recent study, we described a new class of outer-membrane β-barrels composed of more than one barrel domain and surprisingly came across GfcD and its homologs, which formed the largest cluster of proteins in our dataset (7). These proteins were confidently predicted to contain two barrel domains, an N-terminal one of 12 strands and a C-terminal one, where the assignment of the strand number proved surprisingly difficult, leading us to conclude that it had at least 12 strands, but most likely 13 (an unprecedented number for an outer-membrane β-barrel).

Our study was published shortly before the AlphaFold protein structure prediction method became available, so one of the first proteins we submitted to the new server was GfcD. As anticipated by us, GfcD was predicted as a multibarrel, with an N-terminal 12-stranded domain and a C-terminal 13-stranded one, connected by a periplasmic middle domain. The structure of the C-terminal barrel caught our attention, because it showed a 14-stranded topology missing the last strand, resulting in a lateral aperture between the first (β1) and last (β13) strands. This aperture immediately called to mind other proteins of the outer membrane, for example BamA, LptD, PagP, and FadL, which have been proposed to release hydrophobic ligands into the membrane through a lateral gate (12–17). For LptD, lipopolysaccharide (LPS) export via a 26-stranded outer-membrane β-barrel was simulated by molecular dynamics, showing that, although no aperture is visible in the unliganded structure, a lateral aperture can open at a weak spot in the barrel, between strands β1 and β26, allowing LPS to exit laterally into the membrane (18).

These observations led us to consider that GfcD might be the pore responsible for the export of lipid-anchored exopolysaccharides in group 4 capsules. To explore the plausibility of this inference, we used molecular dynamics simulations, in analogy to the studies on LptD. Specifically, we simulated the full, mature GfcD protein embedded in an *E. coli* outer membrane patch, with the CPS anchor, lipid A (19), inserted into the C-terminal barrel lumen. In our unsteered molecular dynamics simulations we consistently observed the tails of lipid A exiting from the lateral aperture in GfcD, but no membrane lipids penetrating the pore. These findings support a role for GfcD in the export of lipid-anchored exopolysaccharides and expand the range of outer-membrane proteins using the lateral-exit mechanism for the export of hydrophobic ligands.

## Methods

### GfcD modeling and system assembly

No experimental structures of GfcD have been determined, so we employed the model from the AlphaFold2.0 protein structure database (https://www.alphafold.ebi.ac.uk/entry/P75882) (20). To obtain the mature form of the protein, we removed the N-terminal signal sequence (residues 1-18) as detected by SignalP (https://services.healthtech.dtu.dk/services/SignalP-5.0/) (21,22). We then built a liganded version of this mature form by inserting a lipid A moiety interactively into the channel of the C-terminal barrel using Pymol, ensuring that the disaccharide head was oriented towards the extracellular side and that there were no major clashes. We refined this starting GfcD-lipid A complex with a vacuum energy minimization, keeping the position of all protein heavy atoms restrained. For simulations, we embedded both the liganded and unliganded mature GfcD forms into a model *E. coli* outer membrane using CHARMM-GUI (23–26). This model outer membrane is composed of (i) an outer leaflet of deep rough lipopolysaccharides (Re LPS), characterized by a minimal core of two 3-deoxy-D-manno-octulosonic acid (KDO) molecules attached to the lipid A and the absence of O-antigen chains, and (ii) an inner leaflet made of a mixture of 1-palmitoyl-2-oleoyl phosphatidylethanolamine (POPE), 1-palmitoyl-2-oleoyl-sn-glycero-3-phosphatidylglycerol (POPG), and 1,1′-palmitoyl-2,2′-vacenoyl cardiolipin (PVCL2) with a 18:1:1 ratio. Such compositions are well-established for the simulation of the *E. coli* outer membrane (27–30). We neutralized the charge on the LPS molecules with Ca2+ ions, and further adjusted the ionic strength of the system with 150 mM NaCl. The complex was solvated with TIP3P waters, resulting in a system of 236 Å x 236 Å x 137 Å dimension, comprising 724742 atoms in the unliganded version, and of 149 Å x 149 Å x 170 Å dimension, comprising 347630 atoms in the liganded version. The assembly procedure is summarized in **Figure S1**.

### Molecular dynamics simulations

All simulations were performed in GROMACS 2020.6 (31–33) with the CHARMM 36m forcefield (34– 36), using the LINCS algorithm to constrain bond vibrations (37). Non-bonded interactions were calculated to a cutoff of 1.2 nm, and van der Waals forces were smoothly decreased to 0 beyond 1 nm distance. Long-range electrostatic interactions were calculated using particle mesh Ewald (38).

We ran one simulation for the unliganded form of GfcD and 4 independent simulations for the liganded form, one of these without the C-terminal plug. In each case, we performed 30.375 ns of equilibration divided into six consecutive steps, where the system was brought to temperature and atomic restraints were gradually released (summarized in Table S1). The production runs were 1-µs long, with a 2 fs timestep. Temperature was held at 303.15 K with the stochastic velocity rescaling thermostat (39), using a coupling constant of 1 ps. Pressure was kept at 1 bar using a semi-isotropic Parrinello-Rahman barostat (40,41), using a time constant of 5 ps in all directions, and compressibility values of 4.5 x 10^-5^ bar^-1^ for XY and Z.

### Trajectory analysis

Calculation of angles, root-mean-square deviation and fluctuations were performed with GROMACS. The lipid rotation angle was monitored with a vector connecting the two phosphorus atoms in the lipid A head (PA, PB) in relation to the Z-axis of the simulation box.

We estimated the insertion of lipid A into the membrane with VMD (42), by computing the fraction of the lipid A area penetrating into the membrane during the simulations.

Other analysis scripts were written in Python using MDAnalysis (43,44).

Molecular graphics have been produced in Pymol, Protein Imager (45), and Biorender (https://www.biorender.com/).

### Sequence analysis and database searches

We performed sequence analyses in the MPI Bioinformatics Toolkit (https://toolkit.tuebingen.mpg.de/) (46,47), using database versions of August 2022. We identified AlphaFold models for GfcD homologs using a BLAST search against the alphafold_uniprot50 database, which gave us access to precomputed AlphaFold models contained in the Uniprot database (https://www.uniprot.org/), filtered to a maximum pairwise sequence identity of 50%.

## Results

### The GfcD C-terminal barrel contains a lateral gate into the membrane

In the absence of experimental GfcD structures, we used an AlphaFold model for biophysical simulations of the protein. This model is composed of two membrane-embedded β-barrels, separated by a periplasmic domain, in close agreement with a previous computational study based on sequence comparisons and coevolution analysis (7). The N-terminal barrel (GfcD-N) is a canonical 12-stranded structure, but the C-terminal one (GfcD-C) is a highly unusual form, which has the topology of a 14-stranded barrel but is composed only of 13 strands. The lack of the 14th strand in this fold results in a lateral opening between strands β1 and β13, which leads directly into the membrane (**Figure 1**). Furthermore, the C-terminal end of the protein is folded into the channel formed by this barrel, in a manner reminiscent of a plug (**Figure 1**). Combined, this led us to the hypothesis that the C-terminal barrel is responsible for the export of lipid-anchored exopolysaccharides, with the lateral pore acting as a gate for the lipid anchor to enter the membrane. To test the reliability of the topology we predicted for GfcD, we extracted all AlphaFold2 models of GfcD homologs from the AlphaFold-Uniprot database and sampled this dataset at sequence identities to GfcD ranging from 45% to 25% (**Figure S2**). Superposition of the mature form of these 5 models with GfcD yielded pairwise root mean-square deviation (RMSD) measurements of between 2.0 and 3.2 Å. Limiting the alignment to the C-terminal barrel yielded even lower pairwise RMSD values, between 0.8 and 1.9 Å (**Figure S2**). Thus, the overall fold is essentially the same in all models. More importantly, the lateral opening between strands β1 and β13 was clearly apparent and of similar dimensions in all models (**Figure S2**).

**Figure 1.**
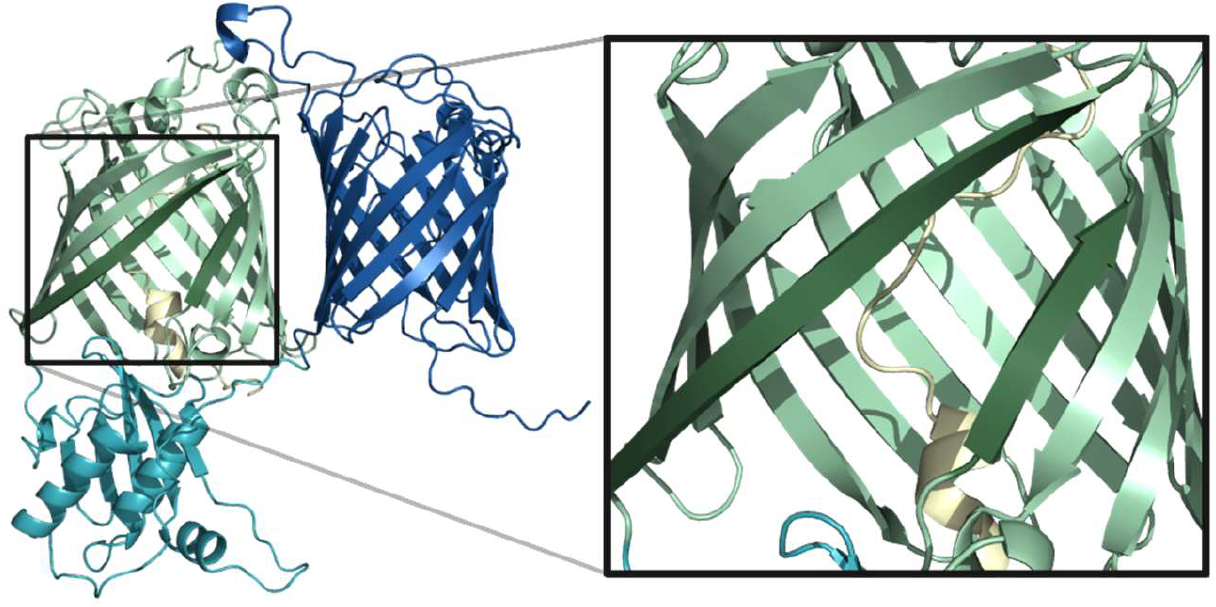
View of GfcD, showing its three domains: GfcD-N (blue; residues 19-278), GfcD-M (cyan; residues 279-423), and GfcD-C (green; residues 424-698), with its C-ter plug highlighted in yellow (residues 674-698). The inset provides a close-up of the lateral gate in GfcD-C.

### The lateral gate in GfcD is stable in molecular dynamics simulations

To test the stability of the predicted GfcD structure we inserted the full, mature protein into a model outer membrane and performed a 1-µs-long molecular dynamics simulation (see Methods). We observed backbone RMSD values of between 2 and 4 Å relative to the starting structure (**Figure 2**), consistent with values reported for other membrane-embedded β-barrel protein simulations (48–53). Backbone root mean-square fluctuation (RMSF) was also low, except for four distinct regions of high flexibility (RMSF > 4 Å): the regions at the N-and C-termini, the extracellular loop around residue 230, and the junctions between the periplasmic domain and the two barrels (**Figure 2**). Overall, the secondary structure and fold of GfcD was stable throughout the simulations, consistent with previous studies of membrane-embedded β-barrels (48,50). Importantly, in the simulations the lateral gate remains unchanged. To further explore the behavior of this region of the protein, we monitored the gate size by measuring the distances between the Cα carbons of three pairs of residues, evenly distributed along the gate (LEU437-TRP668, LEU433-VAL666, THR427-ARG663). For all measured distances, the aperture barely fluctuates. Certainly it does not close (**Figure 2**), further supporting the hypothesis that this represents a bona fide lateral exit into the membrane. Interestingly, while the gate remains open, lipids from the membrane never penetrate into the central channel. Instead, despite its relatively hydrophobic characteristics, the inner channel of the protein is quickly hydrated.

**Figure 2.**
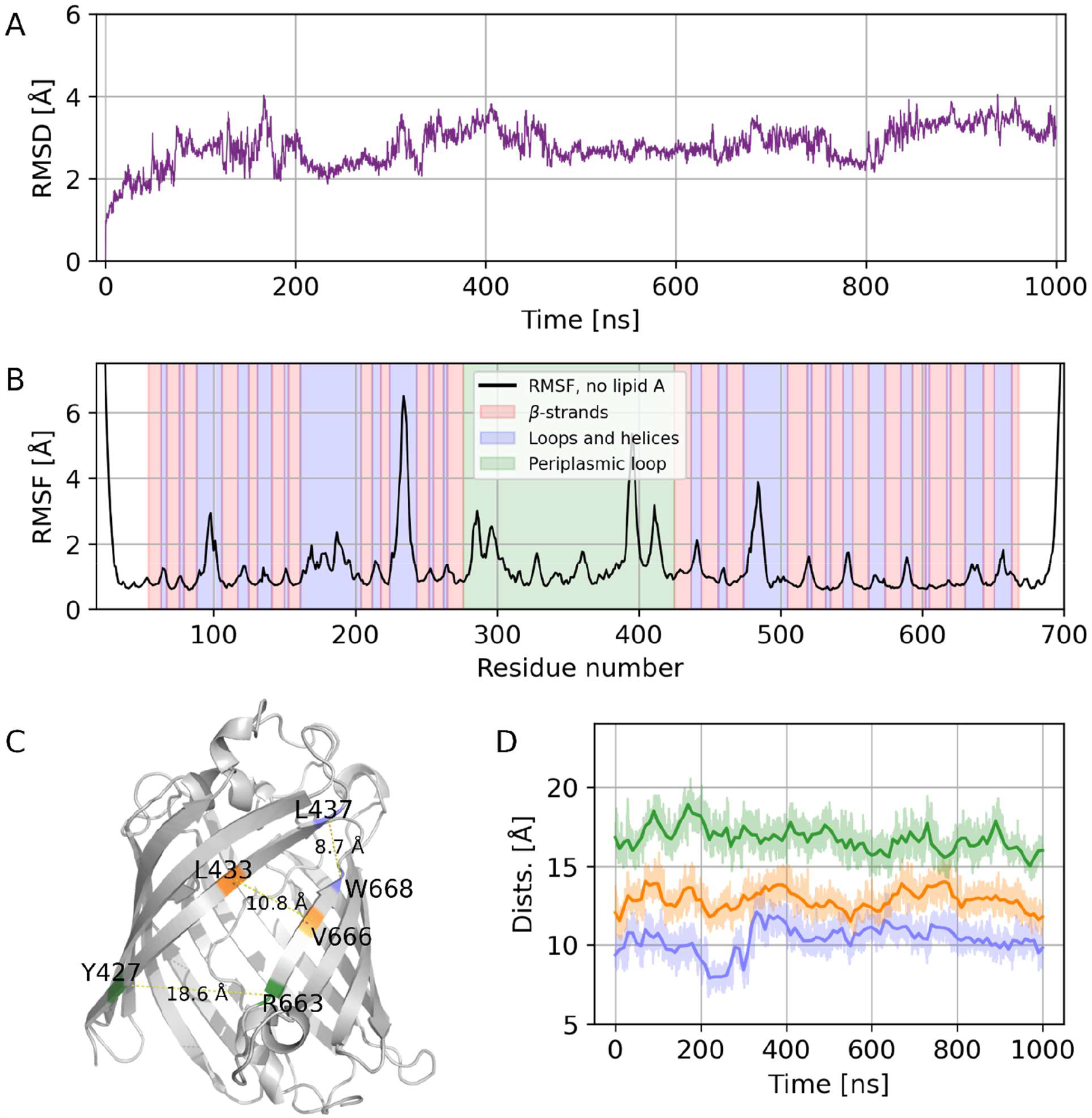
The lateral aperture of GfcD is stable in the unliganded form. (A) RMSD of the protein backbone. (B) RMSF of the protein backbone, with the secondary structure of GfcD highlighted in the background. (C) Three reference distances with which we measure the aperture (L437-W668, L433-W666, Y427-R663). (D) Time evolution of the three distances shown in Panel C.

### GfcD can accommodate lipid A inside the GfcD-C channel without conformational changes

To explore whether the lateral gate can allow the exit of lipid-anchored exopolysaccharides into the outer membrane, we introduced lipid A into the channel of GfcD-C and performed a new set of 1-µs-long molecular dynamics simulations of the full, mature protein. As with the empty GfcD, the protein remained stable during the simulation, with similar backbone RMSD and RMSF values (**Figure 3**). Here as well, the lateral pore remained unchanged throughout the simulation (**Figure 3**). A comparison with the empty structure (**Figure 3E**) shows that the presence of lipid A only results in a slight widening of the periplasmic end of the gate, the rest of the gate remaining the same. Further, the consistently low RMSD values across simulation time show that, even though GfcD is a multidomain protein, these domains do not substantially move relative to each other, whether or not lipid A is present.

**Figure 3.**
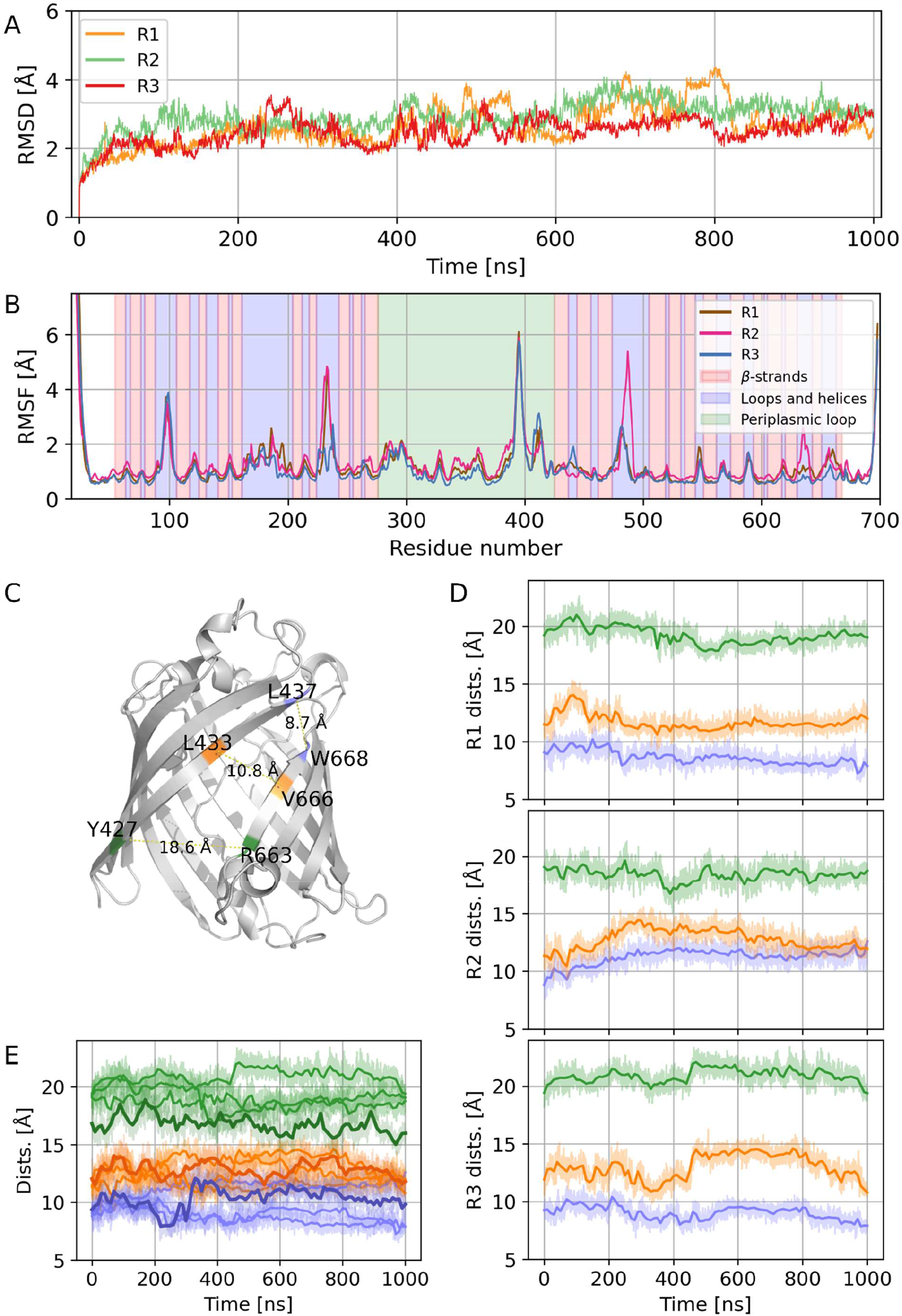
The lateral aperture of GfcD is stable in the simulations with lipid A in the GfcD-C channel. (A) RMSD of the protein backbone. (B) RMSF of the protein backbone, with the secondary structure of GfcD highlighted in the background. (C) Three reference distances with which we measure the aperture (L437-W668, L433-W666, Y427-R663). (D) Time evolution of the three distances shown in Panel C. (E) Aggregated plots for the time evolutions shown in Figure 2D (unliganded) and Figure 3D (with lipid A). To facilitate the comparison, the average lines for the unliganded simulations are shown in bold.

Despite its large size, lipid A can be fully accommodated in the GfcD-C barrel even with the C-terminal extension present in the channel, suggesting that, if this acts as a plug, it does not obstruct the access of lipid A. To test this, we removed the extension (residues ASP674 -GLN698) and ran a further 1-µs-long simulation. As expected, we saw no difference in the behavior of lipid and protein (**Figures S3**,**S4**). A hydration layer is always present between lipid A and the protein, highlighting that the molecule can still move with relative freedom.

To further characterize the behavior of lipid A in the protein channel, we monitored the tilting angle of the molecule by tracking the orientation of its headgroup throughout the simulation time, the lipid tails being too flexible to be used as indicators. We considered the tilting angle relevant for the export process because the gate in GfcD-C is angled relative to the channel, with an inclination set by the shear number of the β-strands in the barrel. While the orientation adopted by lipid A is different in each independent simulation, with values between 112.5 and 180° (**Figure 4**), the orientation becomes relatively stable after 400ns. This variability between replicas is expected, as lipid A should not have a specific binding site within the channel. Most importantly though, despite the considerable span of values, all the angles in the set allowed a rotation of the lipid such that its tails exited the gate.

**Figure 4.**
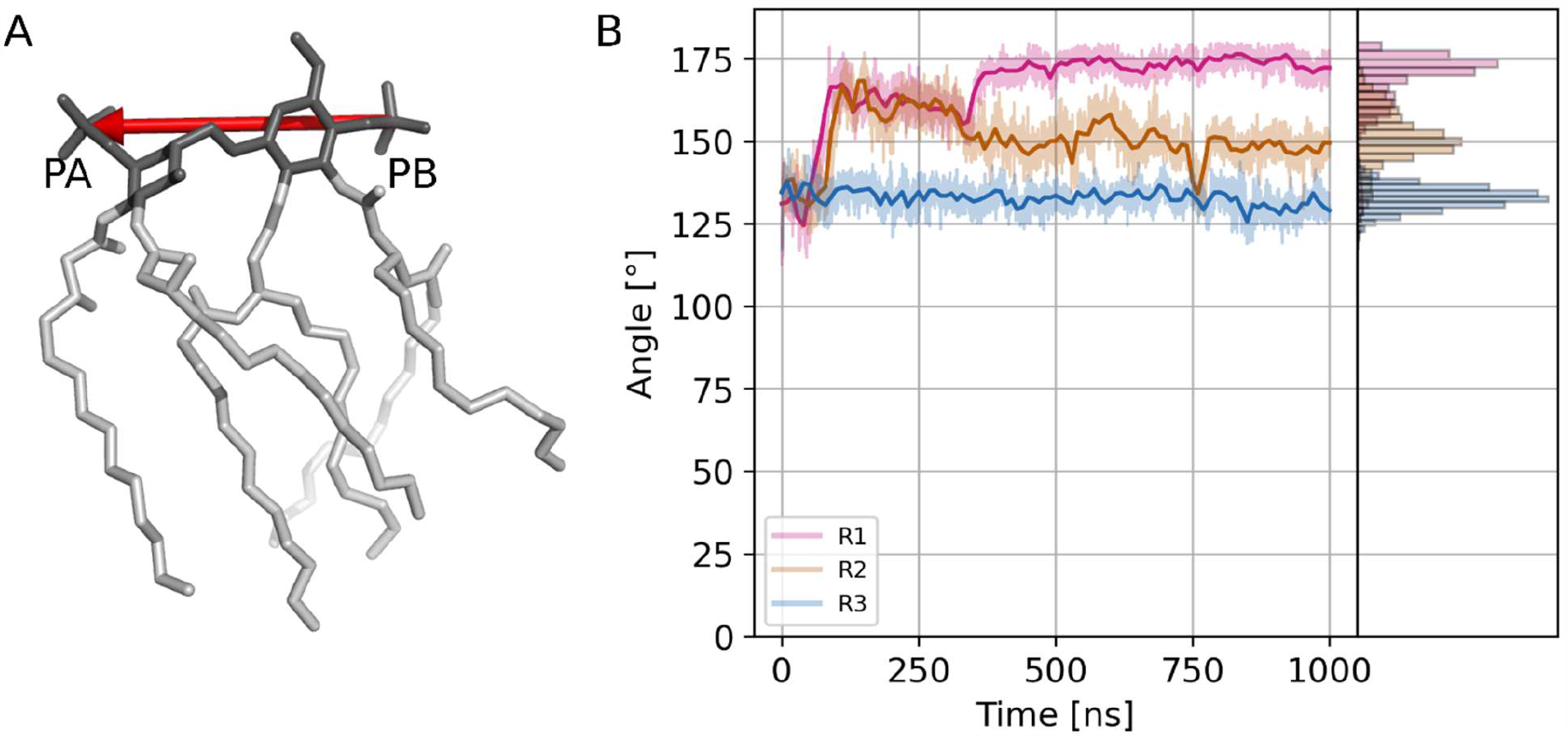
Lipid A spontaneously tilts inside the GfcD-C channel at an angle between 112.5 and 180°. (A) The orientation vector used for monitoring the tilting angle of lipid A during the simulations. The lipid A head is shown in dark grey. (B) Line plot of the tilting angle of lipid A as a function of time, with histograms on the right.

### Lipid A spontaneously crosses the lateral gate in GfcD-C to enter the membrane

Although we simulated lipid A without any biasing force, all our simulations show partial egress of the molecule towards the membrane (**Figure 5, Videos**). To quantitatively assess the exit of lipid A from the barrel, we measured the fraction of lipid A penetrating into the membrane throughout the simulations (**Figure 5**). In all simulations, lipid A rapidly exited the channel to about 20%, in two within the first few nanoseconds, but in the third more gradually to 100 ns. This did not correspond to a partial membrane exposure of all six lipid tails, but rather to the complete exit of one tail, which was the one closest to the PB phosphate group in all three simulations, and the partial exit of the nearest second tail. In the simulation in which the exit proceeded gradually, we observed a retreat of the second tail back into the barrel, followed by its partial reemergence over the next 300 ns, after which the process stalled. In a second simulation, the process also stalled after the initial rapid exit of the lipid tails, but towards the end of the runtime a third lipid tail started to exit, before the second had fully done so. Only in the third simulation did the exit proceed to completion within the runtime, i.e. to the full membrane insertion of all six tails, albeit with occasional partial retreats of tails that had already started to exit.

**Figure 5.**
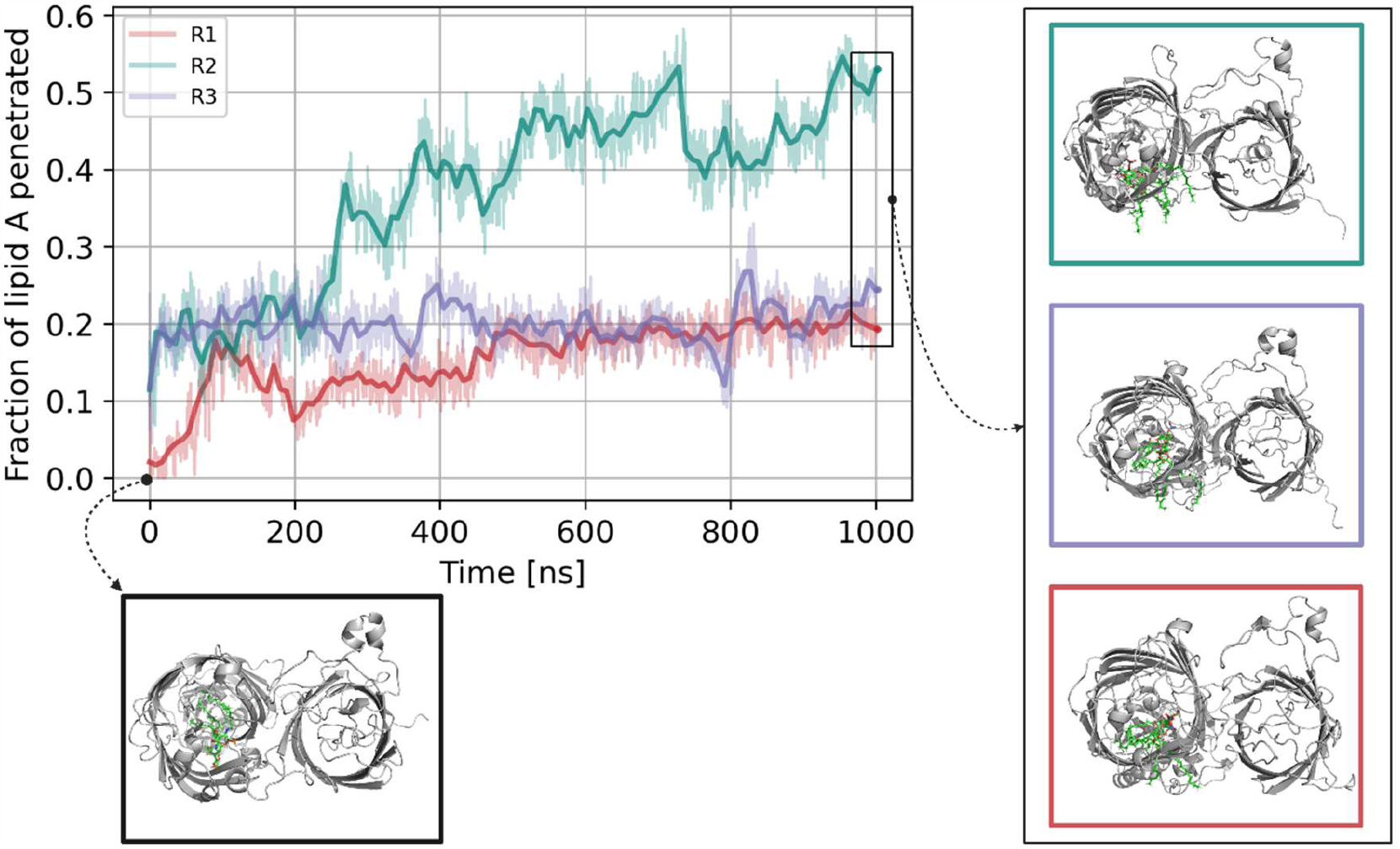
Lipid A exits the GfcD-C channel in all our simulations. The top left panel shows the time evolution of the fraction of lipid A in contact with the membrane. The other four panels show snapshots of the initial and final configurations of GfcD, with lipid A highlighted in green to visualize the egress of its lipid tails. The initial snapshot is framed in black, and the three final snapshots are framed in the color of their corresponding time evolution graph.

In order to establish whether any parts of the channel were required to guide lipid A towards the gate in addition to the residues lining the gate itself, we performed a contact-based analysis of the interactions between lipid A and GfcD residues. We monitored these contacts for each trajectory frame of all three replicas and could thus identify specific residues interacting consistently with lipid A (**Figure 6**). We calculated a contact score for each residue as the geometric average of the number of its contacts with lipid A in each simulation and mapped the result onto the structure of GfcD (**Figure 6**). This analysis established that reproducible contacts with lipid A occur only in strands β1, β2, β12, and β13, which flank the gate, in a loop of the periplasmic domain closest to the GfcD-C channel and in the adjacent part of the C-terminal extension. In the one replica in which the tails of lipid A exited to completion, they even contacted the GfcD-N barrel (**Figure 6**). As expected, these contacts were not observed in the two replicas in which the lipid A tails had only exited partially.

**Figure 6.**
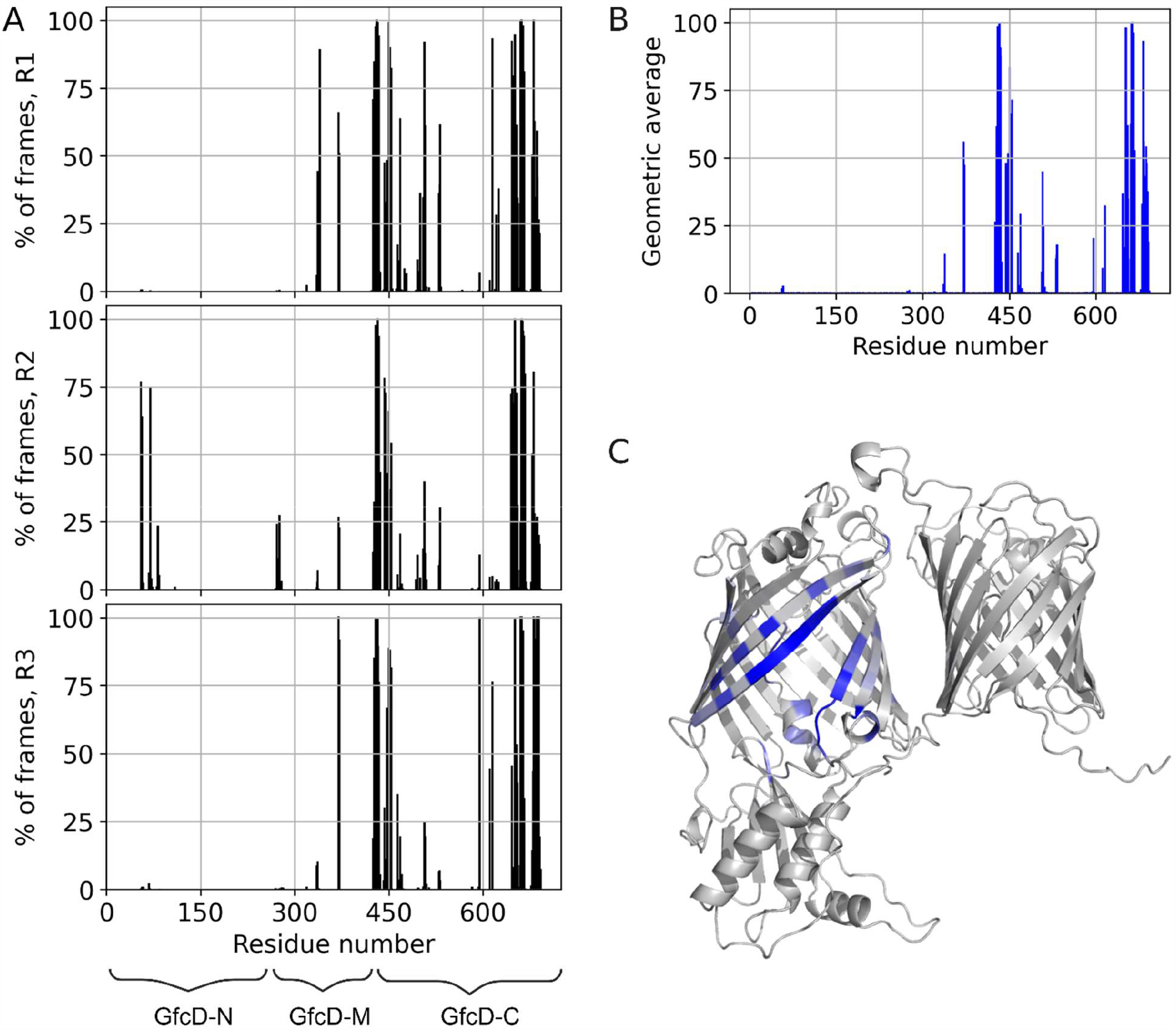
The contacts between GfcD and lipid A characterize the parts of GfcD-C that are directly involved in lipid A export. (A) Percentage of frames during which lipid A was in contact with residues in GfcD, for each replica. (B) The geometric average of the data shown in Panel A. (C) The geometric average mapped onto the structure of GfcD.

## Discussion

In this study, we explored one of the mechanisms by which lipid-anchored capsular polysaccharides might be exported in bacteria with an outer membrane. We found that the GfcD family of proteins provides an attractive candidate for performing this function. GfcD homologs form the largest group of proteins with more than one barrel domain in the outer membrane of Gram-negative bacteria and are present widely in species of this group, tracking the wide-spread occurrence of exopolysaccharide capsules (7). Previous studies have established that *gfcABCD* homologs are involved in exopolysaccharide biosynthesis and transport (3,7–11). Here we used unsteered molecular dynamics simulations to study the full-length, mature GfcD protein of *E. coli* in a model outer membrane patch, with and without lipidA inserted into the GfcD C-terminal barrel. Our results provide further support for an activity of GfcD in the export of lipid-anchored polysaccharides and outline a mechanism for this process.

In our analyses, the lateral gate in the C-terminal barrel of GfcD (GfcD-C) proved to be consistently predicted across the full range of homologs, from nearly identical ones to ones that had diverged into the “midnight zone” of homology below 20% sequence identity (54). This gate was stably open in molecular dynamics simulations, but prevented the penetration of membrane lipids, possibly because of its slanted angle, while allowing the efficient exit of the lipid tails in lipid A.

The middle domain of GfcD, GfcD-M, which is located at the periplasmic entrance of GfcD-C, could be considered a regulator of this exit process. The C-terminal plug of GfcD-C could also be part of a gating mechanism, acting synergistically with GfcD-M to regulate access to the GfcD-C lumen, but we found that this plug does not impede the positioning of the lipid anchor inside the GfcD-C channel and that its removal does not lead to significant changes in the lateral exit of the lipid tails. Its possible role in regulating exopolysaccharide exit thus remains to be elucidated.

Concerning the molecular mechanism underlying the export process, we hypothesize that the saccharide moieties of the exopolysaccharides traverse the channel first and their hydration provides a strong pulling force that drags the whole molecule towards the environment. Once the exopolysaccharide has gone through the channel and lipid A is inserted in GfcD-C, it spontaneously tilts, allowing its hydrophobic tails to exit the channel through the lateral gate into the membrane, in a process similar to that envisaged for LPS export through LptD (13,14,18,55). In our simulations, the tails do not exit the GfcD-C channel simultaneously, but in a sequential process that may stall partway through the process. We envisage that the pulling force exerted by the hydration of the saccharide moieties could be important to overcome such energy barriers during the export process.

Overall, our study provides a viable hypothesis for the biological role of GfcD homologs in the export of lipid-anchored exopolysaccharides, with potentially broader implications for the lateral-exit model of hydrophobic macromolecule export into the bacterial outer membrane through lateral gates in outer-membrane β-barrels.

## Author Contributions

Cecilia Fruet: set up and performed all molecular dynamics simulations, analyzed data, generated the figures, co-wrote the article; Mikel Martinez-Goikoetxea: performed the bioinformatics analyses, advised the data analysis, co-wrote the article; Felipe Merino: advised the molecular dynamics simulations and data analysis, co-wrote the article; Andrei N. Lupas: conceptualized the project, guided the bioinformatic analyses, provided the funds for the project, co-wrote the article. All authors participated in generating the final form of the article.

## Acknowledgments

This work was supported by institutional funds from the Max Planck Society. The MD simulations were performed using the Raven cluster, part of the Max Planck Computing and Data Facility (https://www.mpcdf.mpg.de).

Cecilia Fruet acknowledges Martin Oettel, Raffaello Potestio, and Thomas Tarenzi for helpful and insightful discussions. Cecilia Fruet is grateful to the Universities of Trento and Tübingen for the scholarship awarded for participation in the Double Degree Program between the two Universities.

