## Supplemental Material for "A computational model for lipid-anchored polysaccharide export by the outer-membrane protein GfcD"

**3** Present address: SIB Swiss Institute of Bioinformatics, CH-1015 Lausanne, Switzerland

**4** Present address: Cube Biotech GmbH, D-40789 Monheim, Germany

**Figure S1.** The procedure we used to set up the simulation box for the unliganded and liganded simulations.

### Unliganded version

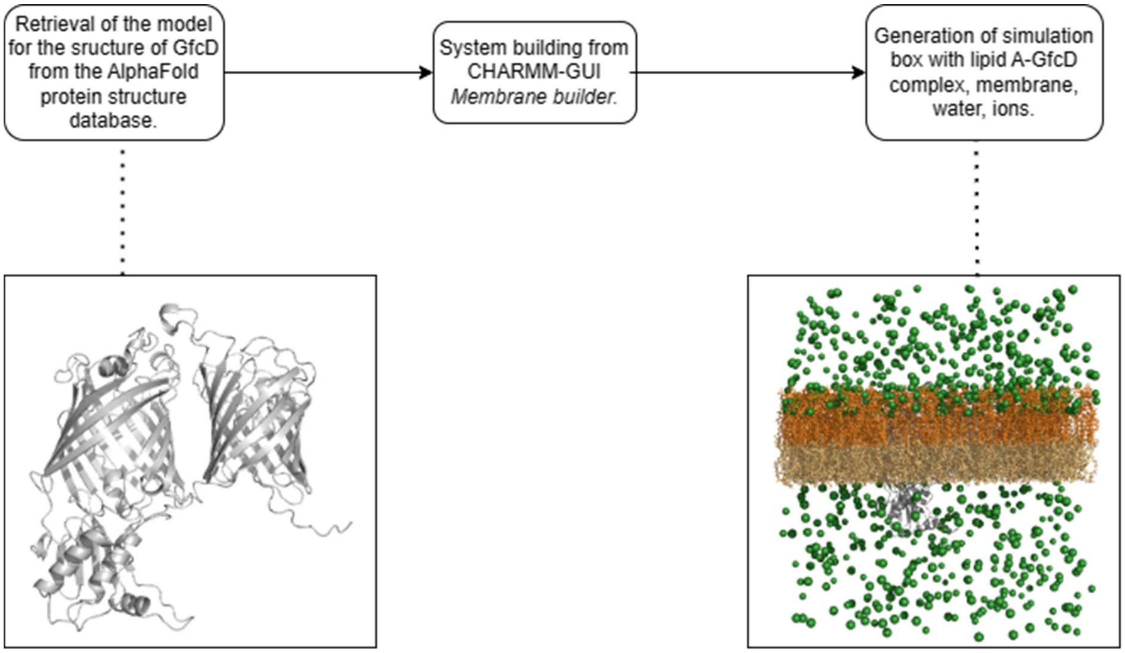


### Liganded version

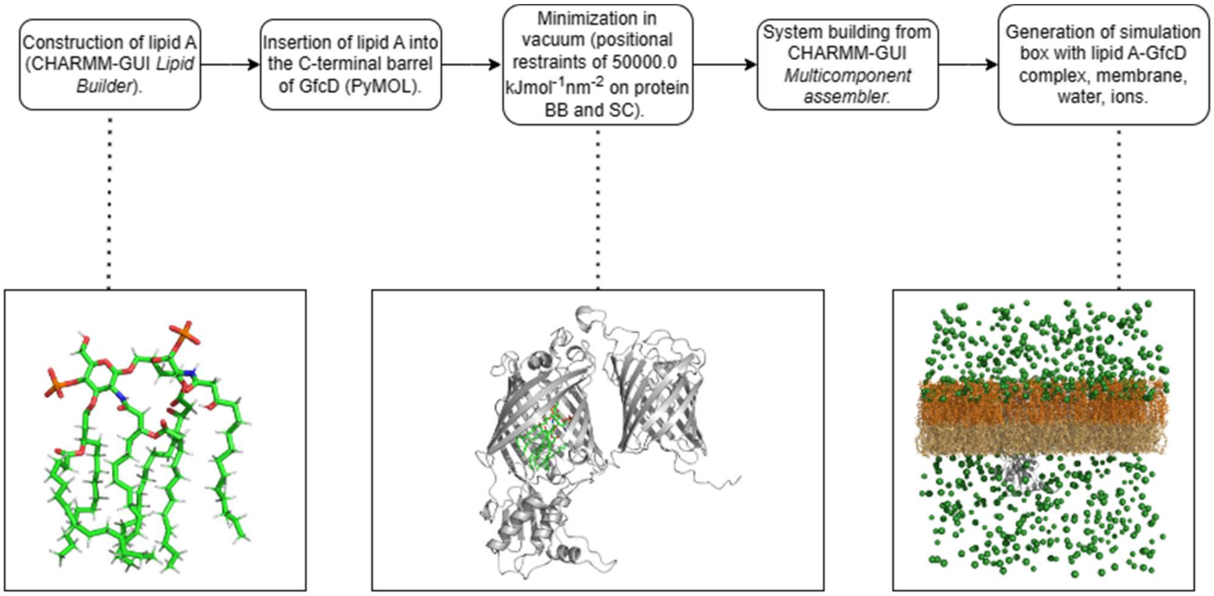

**Table S1.** Summary of the equilibration protocol for our simulations.

| | Equilibration phases | | | | | |
|---|---|---|---|---|---|---|
| | NVT | NVT | NPT | NPT | NPT | NPT |
| Number of steps | $1.25 \times 10^5$ | $1.25 \times 10^5$ | $1.25 \times 10^5$ | $5 \times 10^6$ | $5 \times 10^6$ | $5 \times 10^6$ |
| Timestep [ps] | $1 \times 10^{-3}$ | $1 \times 10^{-3}$ | $1 \times 10^{-3}$ | $2 \times 10^{-3}$ | $2 \times 10^{-3}$ | $2 \times 10^{-3}$ |
| Constraint algorithm | LINCS | LINCS | LINCS | LINCS | LINCS | LINCS |
| Constraints | H-bonds | H-bonds | H-bonds | H-bonds | H-bonds | H-bonds |
| Thermostat | Berendsen | Berendsen | Berendsen | Berendsen | Berendsen | Berendsen |
| Reference temperature [K] | 303.15 | 303.15 | 303.15 | 303.15 | 303.15 | 303.15 |
| Time constant [ps] | 1.0 | 1.0 | 1.0 | 1.0 | 1.0 | 1.0 |
| Barostat | / | / | Berendsen | Berendsen | Berendsen | Berendsen |
| Coupling type | / | / | semi-isotropic | semi-isotropic | semi-isotropic | semi-isotropic |
| Reference pressure [bar] (same in all directions) | / | / | 1.0 | 1.0 | 1.0 | 1.0 |
| Compressibility [$bar^{-1}$] (same in all directions) | / | / | $4.5 \times 10^{-5}$ | $4.5 \times 10^{-5}$ | $4.5 \times 10^{-5}$ | $4.5 \times 10^{-5}$ |
| Time constant [ps] (same in all directions) | / | / | 5.0 | 5.0 | 5.0 | 5.0 |
| Protein backbone restraints [kJ $mol^{-1}$ $nm^{-2}$] | 4000.0 | 2000.0 | 1000.0 | 500.0 | 200.0 | 50.0 |
| Protein sidechains restraints [kJ $mol^{-1}$ $nm^{-2}$] | 2000.0 | 1000.0 | 500.0 | 200.0 | 50.0 | 0.0 |
| Lipid restraints [kJ $mol^{-1}$ $nm^{-2}$] | 1000.0 | 400.0 | 400.0 | 200.0 | 40.0 | 0.0 |
| Dihedral restraints [kJ $mol^{-1}$ $nm^{-2}$] | 1000.0 | 400.0 | 200.0 | 200.0 | 100.0 | 0.0 |

**Table S2.** Summary of the simulations we ran.

| Simulation type | Replicas | Production length |
|---|---|---|
| Membrane + GfcD | 1 | 1 μs |
| Membrane + GfcD + IA | 3 | 1 μs |
| Membrane + GfcD + IA (no C-ter plug) | 1 | 1 μs |

**Figure S2.** (A) Comparison of GfcD to five homologs, selected to span the range of sequence identities to GfcD from 45% to 25%. The proteins are (1) GfcD, (2) UniProt ID A0A7X4W9T3, (3) A0A2N7DHB9, (4) A0A522EX48, (5) A0A2W5NHM9, and (6) A0A2D7DXB3. The pairwise sequence identity matrix was obtained with Clustal Omega (https://www.ebi.ac.uk/Tools/msa/clustalo/). (B) Pairwise Cα-RMSD (left) and superposition (right) for the C-terminal barrels of the proteins in Panel A.

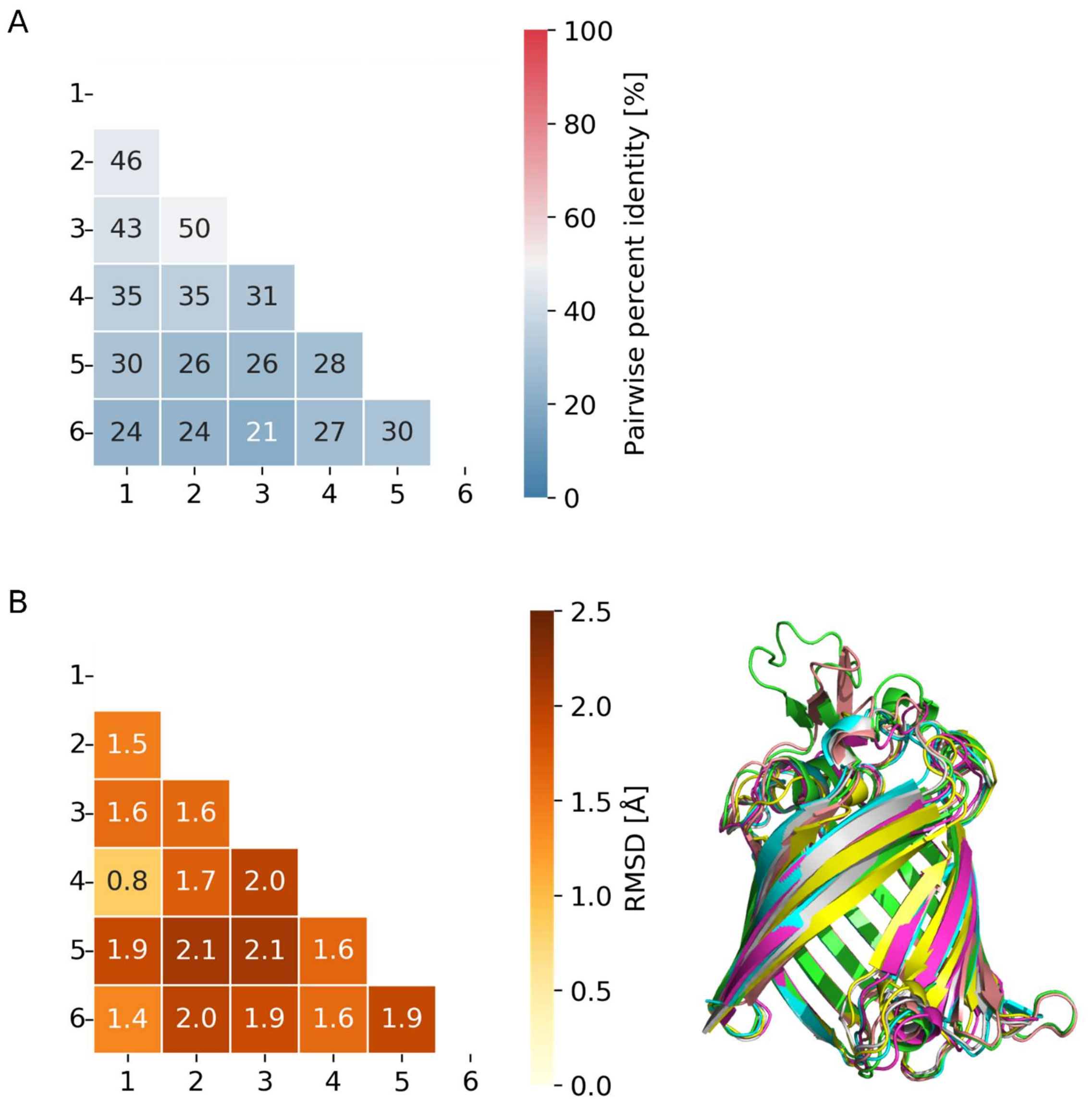

**Figure S3.** The lateral aperture of GfcD is stable in simulations of GfcD without C-terminal plug, with lipid A. (A) RMSD of the protein backbone. (B) RMSF of the protein backbone, with the secondary structure of GfcD highlighted in the background. (C) Three reference distances with which we measure the aperture (L437-W668, L433-W666, Y427-R663). (D) Time evolution of the three distances shown in Panel C.

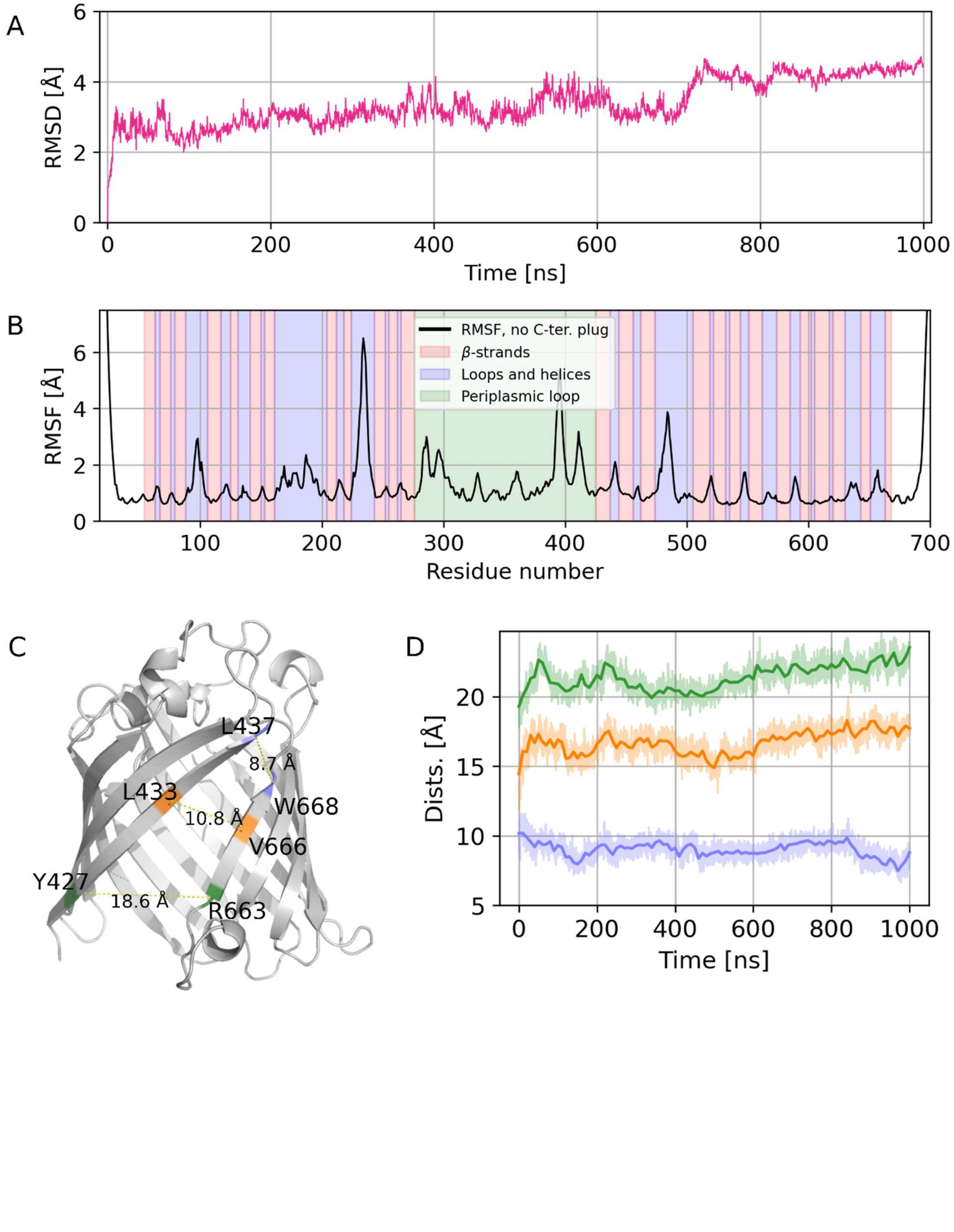

**Figure S4.** Analysis of the simulations of GfcD without C-terminal plug, with lipid A. (A) Tilting angle of lipid A (cf. Figure 4). (B) Insertion of lipid A tails into the membrane (cf. Figure 5).

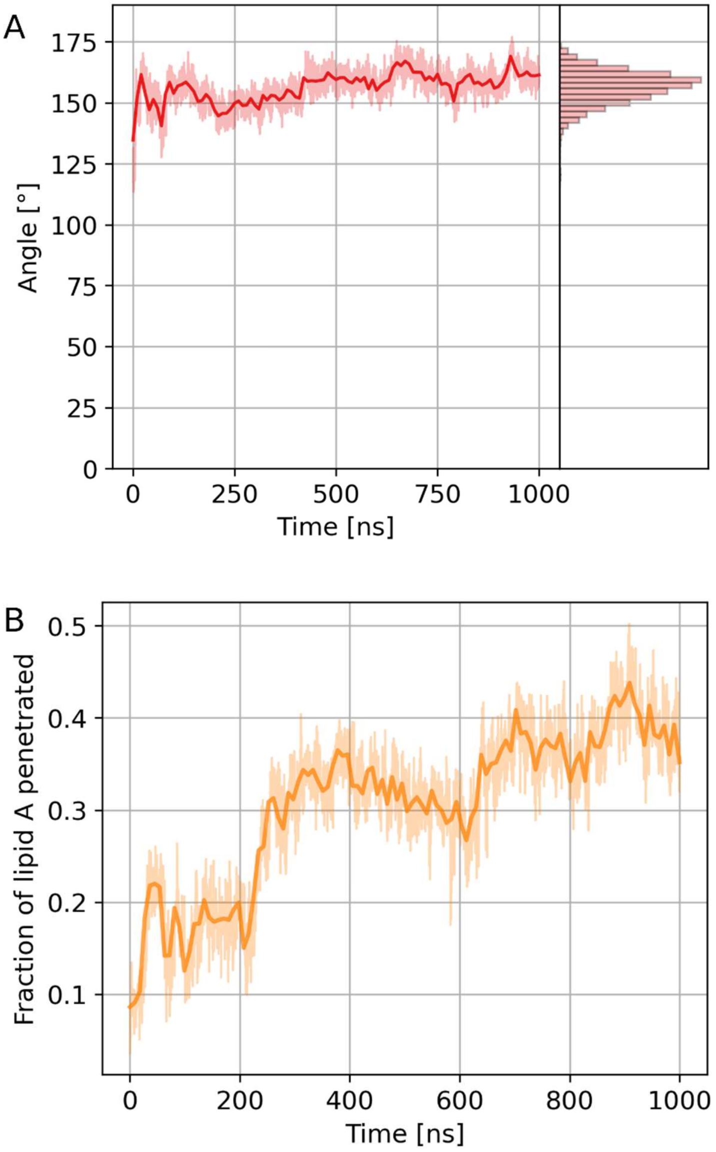
